# IMPACT OF ADOLESCENT INTERMITTENT ETHANOL EXPOSURE IN MALE AND FEMALE RATS ON SOCIAL DRINKING AND NEUROPEPTIDE GENE EXPRESSION

**DOI:** 10.1101/2021.10.29.466460

**Authors:** Trevor T. Towner, Kimberly M. Papastrat, Linda P. Spear, Elena I. Varlinskaya, David F. Werner

## Abstract

**Background:** Alcohol use during adolescence can alter maturational changes that occur in brain regions associated with social and emotional responding. Our previous studies have shown that adult male, but not female rats demonstrate social anxiety-like alterations and enhanced sensitivity to ethanol-induced social facilitation following adolescent intermittent ethanol (AIE) exposure. These consequences of AIE may influence adult social drinking in a sex-specific manner.

**Methods:** To test effects of AIE on social drinking, male and female Sprague-Dawley rats exposed to water or ethanol [0 or 4 g/kg, intragastrically, every other day, between postnatal day (P) 25 and 45] were tested as adults (P72-83) in a social drinking paradigm (30-minute access to a 10% ethanol solution in supersac or supersac alone in groups of three same-sex littermates across two 4-day cycles separated by 4 days off). Social behavior was assessed during the last drinking session, with further assessment of oxytocin (OXT), oxytocin receptor (OXTR), vasopressin (AVP) and vasopressin receptors 1a and 1b (AVPR1a, AVPR1b) in the hypothalamus and lateral septum.

**Results:** Males exposed to AIE consumed more ethanol than water-exposed controls during the second drinking cycle, whereas AIE did not affect supersac intake in males. AIE-exposed females consumed less ethanol and more supersac than water-exposed controls. Water-exposed females drinking ethanol showed more social investigation as well as significantly higher hypothalamic OXTR, AVP, and AVPR1b gene expression than their counterparts ingesting supersac and AIE females drinking ethanol. In males, hypothalamic AVPR1b gene expression was affected by drinking solution, with significantly higher expression evident in males drinking ethanol than those consuming supersac.

**Conclusions:** Collectively, these findings provide new evidence regarding sex-specific effects of AIE on social drinking and suggest that the hypothalamic OXT and AVP systems are implicated in the effects of ingested ethanol on social behavior in a sex- and adolescent exposure-dependent manner.

## Introduction

In the United States, alcohol consumption is normative, with over 50% of people aged 18 years and older reporting drinking alcohol in the previous month (Substance Abuse and Mental Health Services (SAMHSA), 2019). Unfortunately, alcohol use is often initiated during adolescence (Morean et al., 2018), with approximately 2.5 million of 12-17 year old adolescents reporting consuming alcohol for the first time (SAMHSA, 2019). Adolescent alcohol drinking often occurs in social groups (Lipperman-Kreda, Mair, Bersamin, Gruenewald, & Grub, 2015), with common motives for drinking in a social context associated either with the enhancement of positive emotional states or the relief of social anxiety (Blumenthal, Leen-Feldner, Frala, Badour, & Ham, 2010; Kuntsche et al., 2015). During adolescence, the social brain is highly susceptible to early life stressors (Tzanoulinou & Sandi, 2017), and maladaptive choices such as heavy alcohol use may produce long-term detrimental effects on social behavior.

Although adults consume alcohol on more occasions (SAMHSA, 2019), adolescents often report having more drinks per occasion, commonly referred to as binge drinking (4+ drinks for females and 5+ drinks for males during a 2-hour period, resulting in blood ethanol concentrations above 0.08 mg/dL) or even high-intensity binge drinking (10+ drinks in one occasion). For example, approximately 15% of those 18 years of age report binge drinking in the previous two weeks (Patrick & McElrath, 2019), with rates of high-intensity binge drinking at this age over 5% (Patrick & McElrath, 2019). Human studies have identified various short-term behavioral and neural consequences associated with adolescent alcohol consumption (for reviews see Jones, Lueras, & Nagel, 2018; Spear, 2018), however few studies have evaluated the long-term alterations.

The assessment of the long-term consequences of adolescent alcohol use in humans is limited by practical and ethical concerns, whereas the use of preclinical rodent models, allows researchers to evaluate the persistent alterations resulting from alcohol (ethanol) exposure during the adolescent developmental period. Recent studies have identified several long-lasting behavioral consequences of adolescent ethanol exposure (for reviews see Crews et al., 2019; Spear, 2018; Towner & Varlinskaya, 2020). For example, adolescent binge-level ethanol exposure is associated with behavioral flexibility deficits (Coleman, Liu, Oguz, Styner, & Crews, 2014; Fernandez & Savage, 2017; Galaj, Kipp, Floresco, & Savage, 2019; Gass et al., 2014; Varlinskaya, Hosová, Towner, Werner, & Spear, 2020), enhanced anxiety-like behavior (Pandey, Sakharkar, Tang, & Zhang, 2015; Sakharkar et al., 2019; Varlinskaya et al., 2020) and increased risk-taking (Kruse, Schindler, Williams, Weber, & Clark, 2017; Schindler, Tsutsui, & Clark, 2014). In addition, we have consistently identified social behavioral alterations that are evident in adult males, but not their female counterparts, following exposure to ethanol during early-mid adolescence (Dannenhoffer et al., 2018; Varlinskaya, Kim & Spear, 2017; Varlinskaya, Truxell, & Spear, 2014; Varlinskaya et al., 2020). More specifically, following early adolescent intermittent ethanol (AIE) exposure adult males demonstrate reductions in social investigation, play-fighting, and social preference when compared with their water-exposed counterparts.

The observed social alterations following AIE are also accompanied by AIE-associated changes in sensitivity to the social consequences of an acute ethanol challenge. Specifically, males exposed to AIE and challenged with low doses of ethanol in adulthood display increases in social behavior (Varlinskaya et al., 2014). Furthermore, ethanol exposure in adolescence has been shown to impact social behaviors during a social drinking session (Varlinskaya et al., 2017). Males exposed to AIE demonstrated increases in the number of play fighting while drinking socially with cage-mates, an effect of ethanol drinking not observed in water-exposed controls and females (Varlinskaya et al., 2017). Collectively, these studies have identified AIE-induced social impairments accompanied by enhanced sensitivity to the socially facilitating effects of ethanol.

Multiple brain regions including the hypothalamus (HYPO), lateral septum (LS), nucleus accumbens (NAc), amygdala, medial prefrontal cortex (mPFC), and the bed nucleus of the stria terminalis (BNST), are implicated in the regulation of social interactions (for reviews see Ko, 2017; Newman, 1999; Vanderschuren, Achterberg, & Trezza, 2016). Within these regions, the neuropeptides oxytocin (OXT) and arginine vasopressin (AVP), along with their cognate receptors, play substantial roles in modulating different social behaviors (Caldwell, 2017; Johnson & Young, 2017; Veneema & Neumann, 2008; Zoicas, Slattery, & Neumann, 2014) as well as social and non-social anxiety (Bredewold & Veenema, 2018; Dannenhoffer et al., 2018; Harper et al., 2019; Liebsch, Wotjak, Landgraf, & Engelmann, 1996; Lukas et al., 2011). Interestingly, OXT and AVP often have similar effects on social recognition and social memory (Lukas, Toth, Veenema, & Neumann, 2013; Wersinger, Temple, Caldwell, & Young, 2008), while playing opposing roles in the modulation of social anxiety-like behavior. For example, activation of the OXT system increases social approach and social preference (Grund et al., 2019; Lukas et al., 2011), whereas AVP microinjections within the amygdala have been shown to elicit social anxiety-like behavior (Harper et al., 2019).

We have recently found that in male rats, an exposure to a social stimulus increased OXT receptor (OXTR) gene expression in the LS, while decreasing AVP receptor 1a (AVPR1a) gene in the same region (Kim, Varlinskaya, Dannenhoffer, & Spear, 2019). Furthermore, systemic administration of a selective OXTR agonist reverses the social deficits evident in AIE-exposed males (Dannenhoffer et al., 2018). Similarly, administration of a selective AVP receptor 1b (AVPR1b) antagonist was able to restore social behavior in AIE-exposed males, an effect that was not evident with an AVPR1a antagonist (Dannenhoffer et al., 2018). Collectively, these studies support a role of both OXT and AVP peptide systems in social interactions and suggest that a potential dysregulation of these systems might contribute to the social impairments associated with adolescent ethanol exposure.

Previous studies have found that adolescent ethanol exposure promotes greater ethanol consumption in adulthood (for review see Towner & Varlinskaya, 2020), findings that are often reported in isolate-housed animals (Crabbe, Harris, & Koob, 2011). Alternatively, the evaluation of AIE-induced alterations in ethanol consumption under social test circumstances has received limited attention. With the relief of social anxiety often reported as a motive for consuming ethanol in humans, and AIE leading to social anxiety-like behavioral alterations, it is possible that AIE also promotes greater ethanol consumption in social contexts. Furthermore, both OXT and AVP peptide systems are reported to influence ethanol intake (Bahi, Mansouri, & Maamari, 2016; Edwards, Guerrero, Ghoneim, Roberts, & Koob, 2012; Peters, Bowen, Bohrer, McGregor, & Neumann, 2017; Sanbe et al., 2008). For example, ethanol intake was reduced following systemic OXT administration (MacFadyen et al., 2016) as well as systemic administration of an AVPR1b antagonist (Zhou, Rubinstein, Low, & Kreek, 2017; however, see Caldwell et al., 2006). In addition, ethanol exposure and consumption have been shown to alter gene (Allen et al., 2016; Przybycien-Szymanska, Rao, & Pak, 2010; Rivier & Lee, 1996; Silva, Paula-Barbosa, & Madeira, 2002) and protein (Dannenhoffer et al., 2018; Madeira & Paula-Barbosa, 1999; Silva, Madeira, Ruela, & Paula-Basbosa, 2002; Stevenson et al., 2017) expression of these two systems. Together with our previous results that demonstrated AIE-induced alterations of the OXT and AVP systems in the HYPO (Dannenhoffer et al., 2018), it is possible that ethanol ingested under social circumstances might differentially affect these neuropeptide systems in AIE-exposed animals and their water-exposed counterparts.

Given the importance of social motives for drinking in humans and our previous findings of AIE-induced social anxiety-like alterations in male rats, the present study was designed to test the hypothesis that AIE exposure enhances ethanol consumption in a social drinking model in males but not females, with this social drinking also producing social facilitation in AIE-exposed male subjects. In addition, we were interested in determining the effects of social drinking on the OXT and AVP systems in AIE- and water-exposed males and females, therefore expression of genes related to these neuropeptide systems was assessed following the final drinking session in the HYPO and LS. To control for the possibility of ethanol intake being driven by differences in the palatability of the sweetened ethanol solution, intake of the sweetened solution alone was assessed in AIE- and water-exposed males and females.

## Materials and Methods

### Subjects

Male and female Sprague-Dawley rats (N = 192) that were bred and reared in our colony at Binghamton University were used. Animals were housed and maintained on a 12 hour on/off light schedule (lights on at 7:00 AM), in temperature-(21-23 °C) and humidity-controlled vivaria. *Ad libitum* food (Purina Rat Chow, Lowell, Massachusetts) and water were available at all times except during experimental testing. On postnatal day (P) 1, litters were preferentially culled to six males and four females (10 per litter) when possible. On P21, rats were weaned, and group housed in cages of four same sex littermates. Prior to ethanol exposure, all rats were left undisturbed other than routine cage maintenance. All experimental testing was conducted between 15:00 and 17:00 hours under dim light (∼15 lux). The treatment, experimental manipulations, and maintenance of rats was in accordance with the National Institutes of Health guidelines for animal care using protocols approved by Binghamton University Institutional Animal Care and Use Committee.

### Adolescent Intermittent Ethanol (AIE) Exposure

Rats were exposed to ethanol (4 g/kg, 25% v/v) during early-mid adolescence (P25-45) via intragastric gavage every other day, total of 11 exposures. Control rats were exposed to tap water using this same regimen. Animals housed together were assigned to the same adolescent exposure condition. Our previous work has found that this model of ethanol exposure results in blood ethanol concentrations (BECs) between 125 and 200 mg/dL (Kim et al., 2019). After the final ethanol or water exposure (P45), all rats were undisturbed until adulthood (P72) prior to behavioral testing.

### Social Drinking Procedure

Starting on P72, rats were transported to a testing room and placed into a novel cage in groups of three littermates. Following placement, rats were given 30-min access to two bottles containing either 10% ethanol in supersaccharin (supersac: 3% sucrose + 0.125% saccharin; Ji, Gilpin, Richardson, Rivier, & Koob, 2008) or supersac alone, similar to our previous studies (Varlinskaya et al., 2017; Varlinskaya et al., 2015). Two bottles of the same solution were used to eliminate competition at the bottles. A total of eight drinking sessions were given in two 4-day cycles separated by 4 days off. Animals within a cage were given unique markings for later identification. All drinking sessions were video recorded, with time spent drinking scored by a trained experimenter that was blinded to adolescent exposure condition. Drinking was characterized by repetitive tongue movements at the bottle tip, often accompanied by a grasping of the drinking tube with the forepaws. Instances of less than a full second of time at the bottle tip were considered investigation and thus not counted toward drinking time. To calculate individual intake per animal, we first calculated a total intake per second for the cage and multiplied this by the time spent drinking for an individual rat within the cage, equaling the individual intake of solution. Animals were not water restricted prior to the drinking sessions.

During the last drinking test, social investigation (sniffing of cage-mates) and play fighting (number of playful nape attacks) were scored for each individual animal in a drinking group.

### Blood Ethanol Concentration (BEC)

Immediately following the final 30-min drinking session, all animals were rapidly decapitated and trunk blood samples were collected into 1.7 ml tubes. Whole blood was maintained at -80 °C until the time of assay. Aliquots of 25 μl were analyzed using a Perkin Elmer (Waltham, MA, USA) gas chromatograph (Clarus 580) paired with a Perkin Elmer headspace sampler (TurboMatrix 40). Sampling accuracy was confirmed by testing samples of known ethanol concentrations prior to analyzing experimental samples.

### Corticosterone Determination

Whole blood samples were centrifuged at 4°C for 20 minutes at 3000 rpm. Plasma was collected and maintained at -80°C until processing. Male samples were diluted 1:40 and female samples were diluted 1:120 followed by heat-inactivation through immersion in 75°C water for 60-minutes in order to break down endogenous corticosteroid-binding globulin. Samples were then analyzed for corticosterone (CORT) using commercially available ELISA kits (Enzo Life Sciences, Farmingdale, NY, USA). Following heat-inactivation, all samples were treated and processed in accordance with the directions provided by the ELISA kits.

### Tissue Collection and Processing for PCR

Immediately following the last 30-min social drinking session, animals were rapidly decapitated with brains extracted, flash-frozen on dry ice, and stored at -80°C. Using a cryostat (Leica CM 1850, Wetzlar, Germany), brains were sectioned, and regions of interest were collected using tissue micropunches (1.0 - 2.0mm) and in relation to the Paxinos and Waston (2013) brain atlas. Regions of interest collected were the LS and HYPO. Bilateral tissue punches were obtained from each region and stored at -80°C until processing.

Using a Qiagen TissueLyser (Qiagen, Valencia, CA, USA), tissue samples were homogenized through addition of 500 μL of trizol, a 5mm stainless steel bead, and rapidly shaken for 2 minutes. We then added 100 μL of chloroform to each sample and centrifuged at 4°C for 15 minutes. Approximately 250 μL of supernatant was recovered and an equal volume of 70% ethanol was then added prior to being placed in RNAeasy columns (Qiagen, Valencia, CA, USA). Columns were washed using multiple buffers and in accordance with the manufacturer’s instructions. Total RNA was eluded from the column using 30 μL of RNase free water and stored at -80°C. RNA yield and quality were determined using a spectrophotometer (NanoDrop 2000, Thermo Fisher Scientific, Wilmington, DE, USA). Between 0.1 and 1.0 μg (region dependent) of normalized sample were used for cDNA synthesis using QuantiTect Reverse Transcription Kits (Qiagen, Valencia, CA, USA).

Probed cDNA amplification was conducted using 5 μL of SYBR Green Supermix (BioRad, Hercules, CA, USA), 0.5 μL primer, 0.5 μL cDNA sample, and 4 μL of RNase free water for a total reaction volume of 10 μL. Samples were analyzed in triplicate using a 384 well plate and captured in real-time using a PCR detection system (BioRad CFX384).

Cyclophilin A was used as our stable reference gene and the relative expression of the target genes was calculated using the delta-delta (2^-ΔΔC(t)^) method (Livak & Schmitgen, 2001) with water-exposed animals used as the ultimate control group within each sex. The genes of interest and the designed primers are described in Table 1.

**Table 1.**
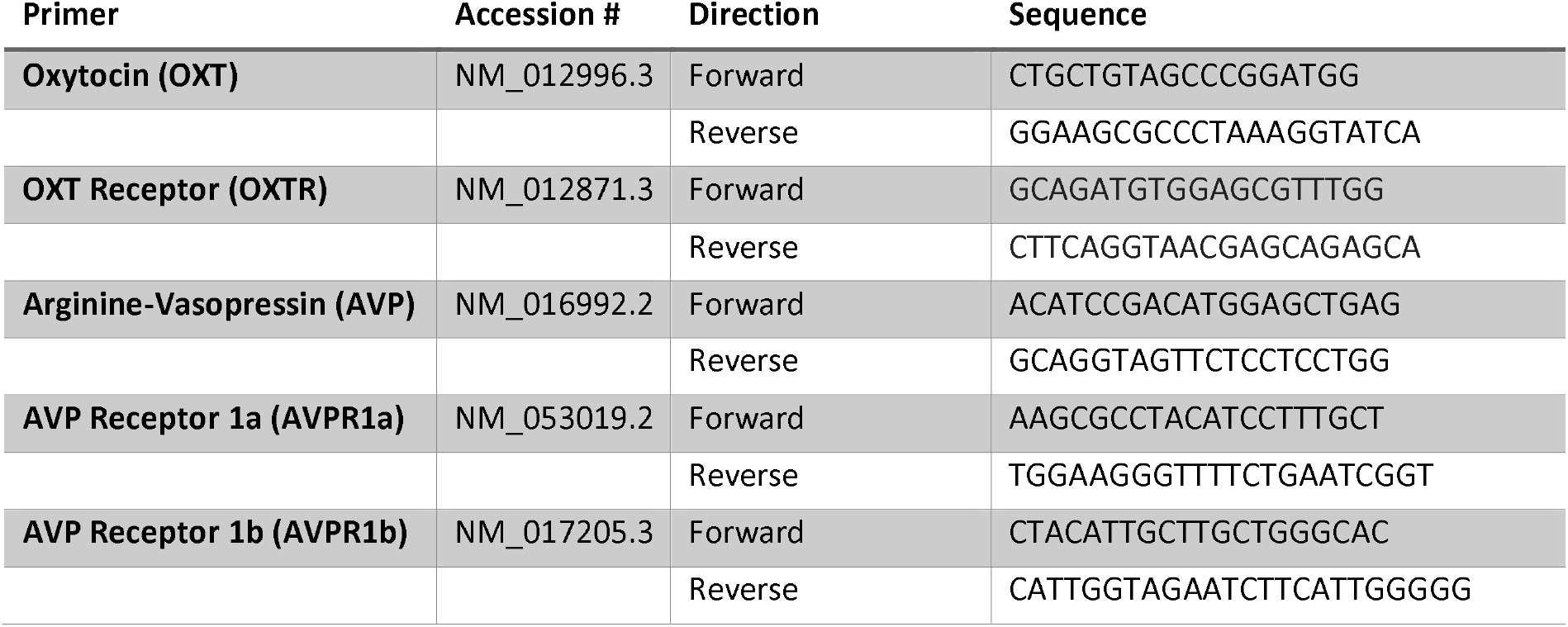
Primers used for real time RT-PCR

### Data Analysis

Ethanol (g/kg) and “supersac” (ml/kg) intake during the social drinking sessions were analyzed using separate 2 (adolescent exposure: water, ethanol) x 2 (sex: male, female) x 8 (drinking test day) repeated measures analyses of variance (ANOVAs) followed by separate for each sex 2-way ANOVAs. BECs assessed on the last drinking day were analyzed using a 2 (adolescent exposure) x 2 (sex) ANOVA. Corticosterone, social investigation, and play fighting were assessed using separate 2 (adolescent exposure) x 2 (drinking solution: ethanol, supersac) x 2 (sex) ANOVAs. When these three-way ANOVAs showed significant interactions involving sex, separate for each 2 (adolescent exposure) x 2 (drinking solution) ANOVAs were performed. Given pronounced sex differences in the OXT/AVP systems at the level of ligands and receptors (Dumais & Veenema, 2016; Rhodes & Rubin, 1999), gene expression data were analyzed separately in males and females using a 2 (adolescent exposure) x 2 (drinking solution) ANOVAs, with the water-exposed group drinking supersac used as ultimate control. Fisher’s planned pairwise comparisons were used to explore significant effects and interactions.

## Results

### Ethanol Intake

A three-way repeated measures ANOVA of ethanol intake revealed main effects of adolescent exposure, F (1, 92) = 8.88, p < 0.005, sex, F (1, 92) = 11.87, p < 0.00, and drinking test day, F (7, 644) = 23.08, p < 0.0001, as well as exposure by sex interaction, F (1, 92) = 12.23, p<0.01. When collapsed across test days, sex differences in ethanol intake were evident in water-exposed controls, with females drinking significantly more than males, whereas AIE-exposed males and females demonstrated similar levels of ethanol intake. Adolescent exposure affected ethanol intake in females, but not males, with AIE females ingesting less ethanol than their water-exposed counterparts (Figure 1A). The ANOVA of ethanol intake also revealed a three-way interaction, F (7, 644) = 2.69, p < 0.01. Further analyses of drinking patterns were conducted separately for females and males. In males, a two-way repeated measures ANOVA revealed a main effect of drinking test day, F (7, 322) = 12.81, p < 0.0001, and adolescent exposure by test day interaction, F (7, 322) = 6.21, p < 0.0001 (see Figure 1B). Post hoc comparisons showed that male subjects significantly increased ethanol intake on test day 2 relative to test day 1 regardless of adolescent exposure condition (ps < 0.05), with AIE males demonstrating relatively stable intake thereafter. In contrast, water-exposed males showed fluctuations in ethanol intake, with significant decreases in intake evident from test day 2 to test day 3 and from test day 5 to test day 6 (ps < 0.05). Furthermore, only water-exposed males substantially increased their intake when tested after a 4-day off period, as evidenced by a significant difference between drinking test days 4 and 5 (p<0.05). Post hoc comparisons of ethanol intake between water- and ethanol-exposed rats revealed significant differences at drinking tests 1, 2, 6, and 8 (ps < 0.05). On test days 1 and 2, water-exposed males consumed more ethanol than AIE males, however AIE males ingested more ethanol on test days 6 and 8 relative to their water-exposed counterparts (see Figure 1B). Further analysis of the cumulative ethanol intake during the first and the second drinking test cycles (test 1 to 4 versus tests 5 to 8, see inserts in Figure 1B) revealed an interaction of drinking cycle and adolescent exposure, F (1, 46) = 14.12, p < 0.001, with AIE males ingesting significantly more ethanol than water-exposed controls during the second, but not the first drinking cycle (p < 0.05).

**Figure 1.**
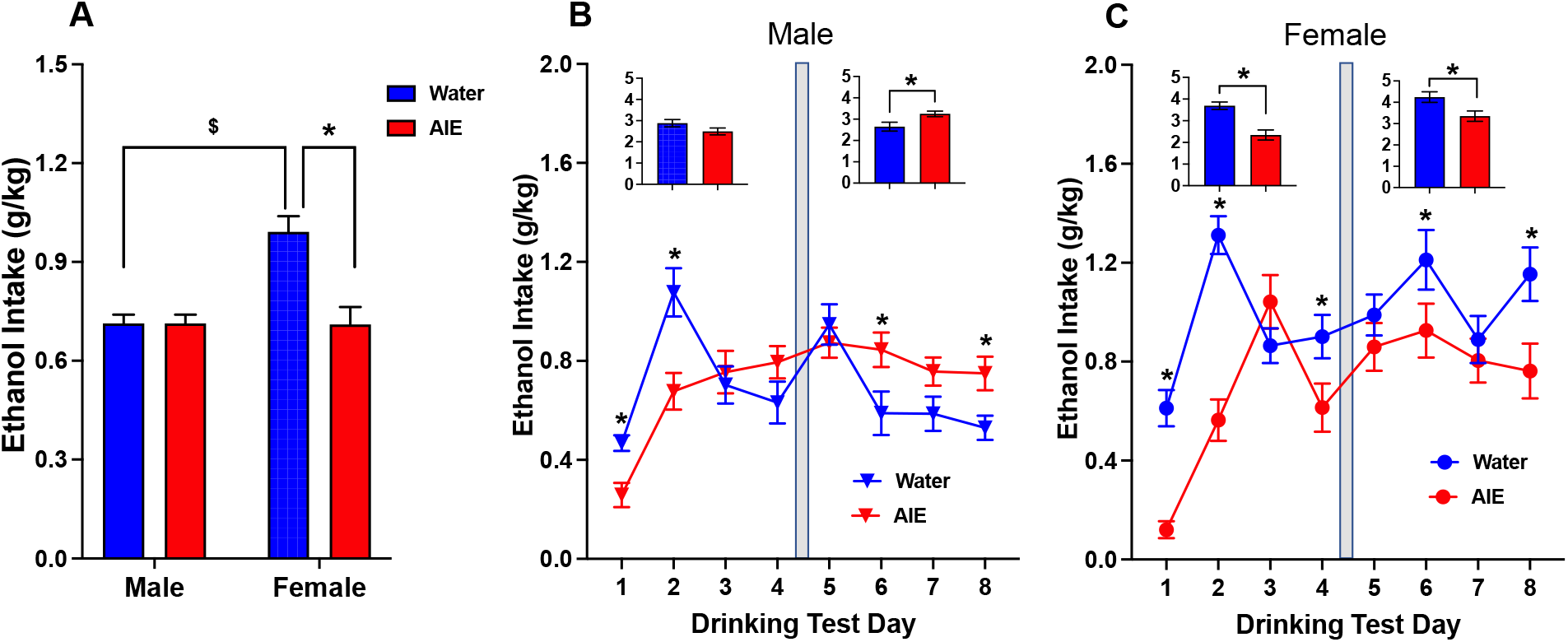
Intake of 10% ethanol in supersac under social drinking circumstances in male and female rats with a prior history of water or AIE exposure. (A) Ethanol intake averaged across drinking test days. Ethanol intake in male (B) and (C) female adult rats during each drinking session. Inserts in B and C represent cumulative intakes during the first and the second drinking cycles of 4 test days, with a 4-day interval between them (marked with a vertical grey bar). A significant difference between males and females in ethanol intake averaged across drinking test days (A) within the same adolescent exposure condition is depicted with $, whereas asterisks (B, C) indicate significant differences in ethanol intake between water- and AIE-exposed animals evident at a certain test day (p < 0.05).

In general, AIE-exposed females consumed significantly less ethanol than their water-exposed counterparts, as evidence by a significant main effect of adolescent exposure, F (1, 46) = 15.71, p < 0.001. A two-way repeated measures ANOVA of female social drinking also revealed a significant main effect of drinking test day, F (7, 322) = 13.52, p < 0.0001, and adolescent exposure by test day interaction, F (7, 322) = 5.67, p < 0.0001. Similar to males, females in both adolescent exposure conditions significantly increased their intake on the second test day relative to the initial social drinking session, with further significant increase evident in AIE females on drinking day 3 (ps < 0.05). In contrast, water-exposed females demonstrated a significant drastic decrease in ethanol intake on day 3 relative to the previous day (p <0.05). Comparisons of AIE and water-exposed conditions on each test day revealed that AIE females had significantly lower intake than their water-exposed counterparts (see Figure 1C) on test days 1, 2, 4, 6, and 8 (ps < 0.05). The cumulative intake in females differed as a function of adolescent exposure regardless of drinking test cycle, F (1, 46) = 15.71, p < 0.001, with AIE females demonstrating significantly lower cumulative intakes during the first (p < 0.0001) as well as the second drinking cycle (p < 0.01) relative to water-exposed controls (see inserts in Figure 1C).

BECs assessed on the last drinking day were significantly correlated with ethanol intake in all experimental groups (Figure 2A-D). A two-way ANOVA revealed a significant interaction of sex and adolescent exposure, F (1,91) = 4.66, p < 0.05, with water-exposed females demonstrating significantly higher BECs than their male counterparts (Figure 2E). BECs in AIE animals, however, did not differ as a function of sex, with no differences in BECs evident between AIE and water-exposed animals regardless of sex.

**Figure 2.**
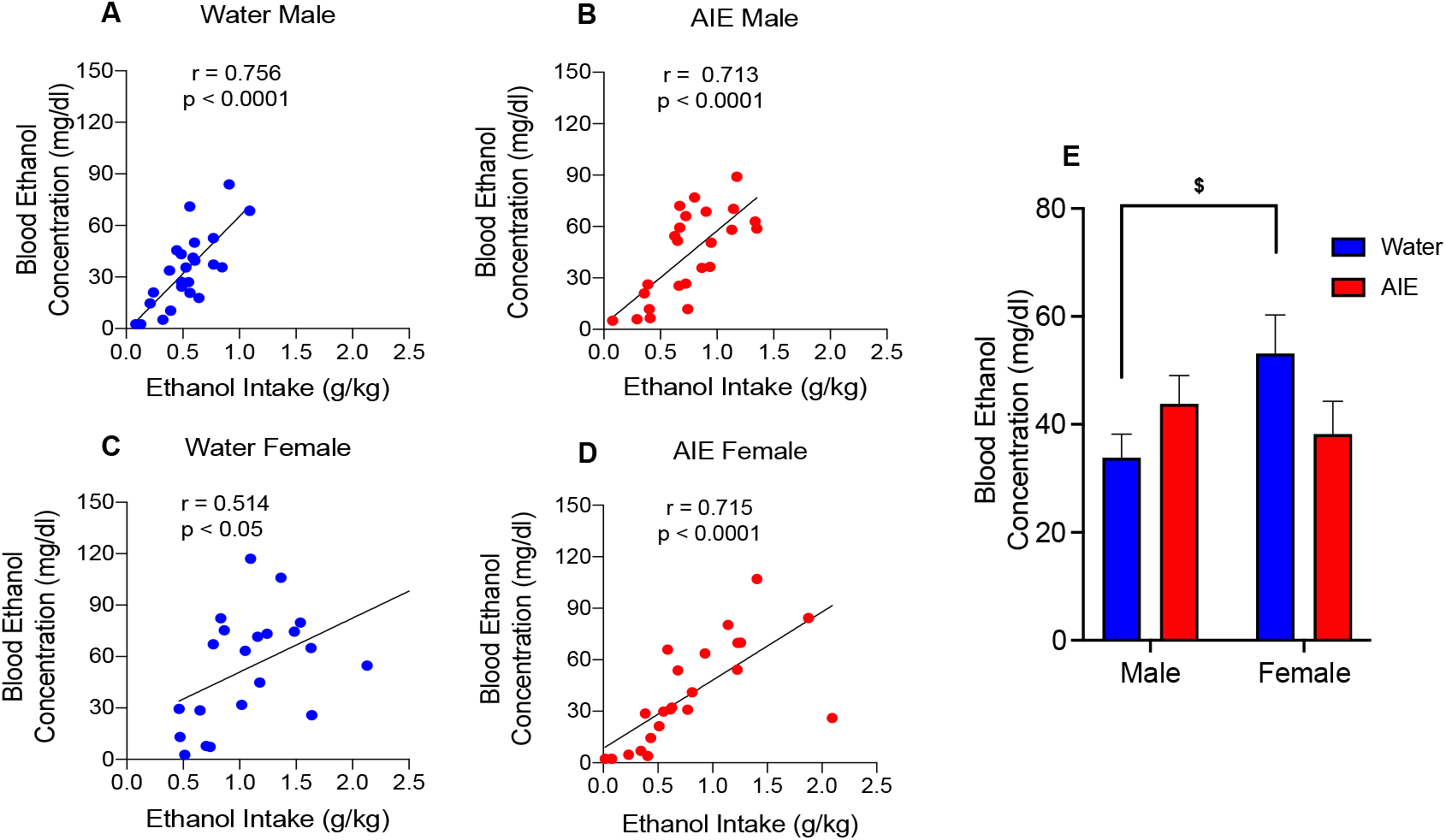
Correlations between ethanol intake and blood ethanol concentrations (BECs) assessed on the last drinking day in water-(A) and AIE-exposed (B) males as well as females exposed to either water (C) or AIE (D). (E) BECs assessed on the last drinking day in males and females. A significant difference between males and females in BECs within the same adolescent exposure condition is depicted with % (p < 0.05).

### Supersac Intake

Since a sweetened with supersac ethanol solution was used for assessment of AIE effects on social drinking, intake of the sweetened solution alone (no ethanol) was assessed in separate cohorts of AIE- and water-exposed males and females in order to determine whether the observed impacts of AIE on social drinking (see Figure 1) were specific to ethanol. A three-way repeated measures ANOVA of supersac intake revealed a main effect of sex F (1, 28) = 52.91, p < .0001 (Figure 3A), with females ingesting significantly more supersac on a mL/kg basis than males. In males, the two-way ANOVA of supersac intake revealed a main effect of test day, F (7, 98) = 33.98, p < 0.0001, with significant increases in supersac intake evident from drinking test day 1 to test day 2 and from test day 4 to test day 5, with no effect of adolescent exposure (Figure 4B). In females, a two-way ANOVA of supersac intake revealed a main effect of test day, F (7, 98) = 39.3, p < 0.0001, and adolescent exposure by test day interaction, F (7, 98) = 2.31, p < 0.05. Females in both adolescent conditions significantly increased supersac consumption from the initial test day to test day 2, with AIE females further increasing intake from day 2 to day 3 and water-exposed females from test day 4 to test day 5 (Figure 3C). In addition, AIE females ingested significantly more supersac than their water-exposed counterparts on drinking test days 4, 6, 7, and 8.

**Figure 3.**
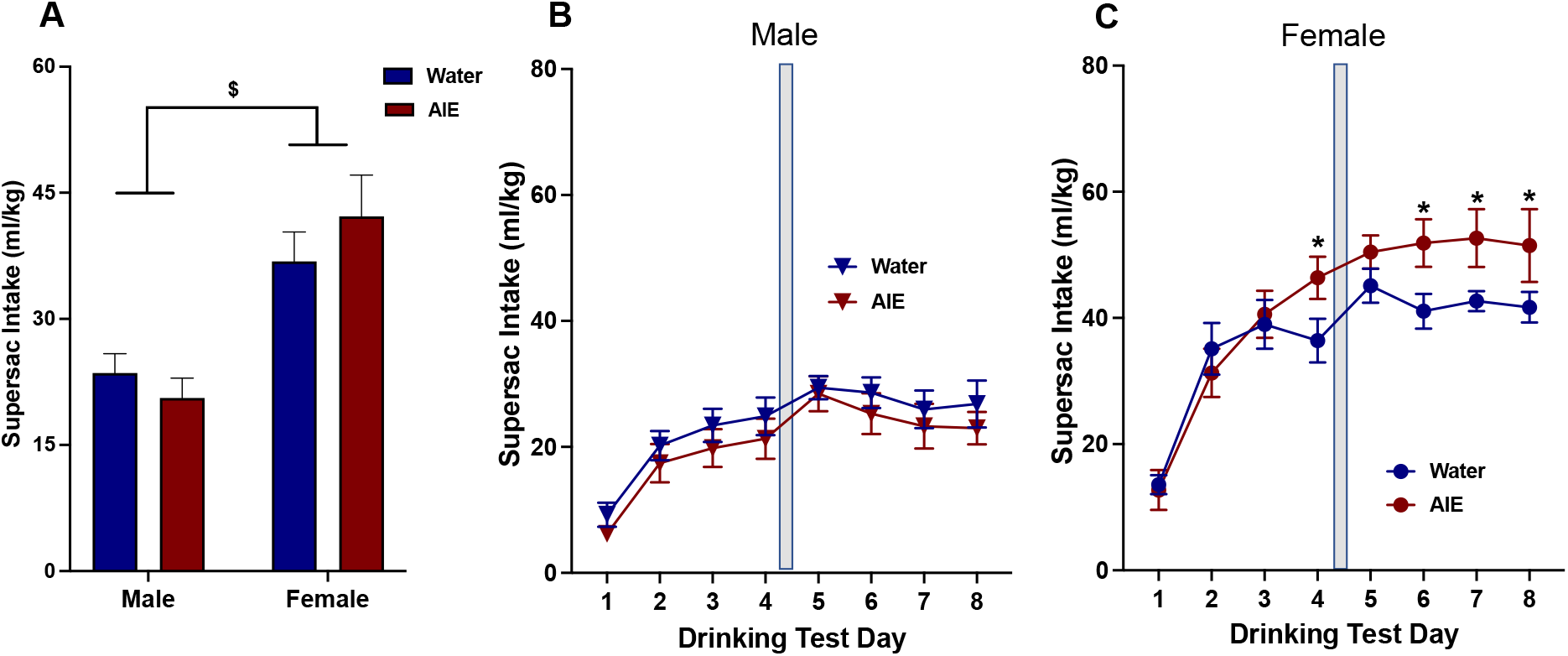
Intake of 10% supersac under social drinking circumstances in male and female rats with a prior history of water and AIE exposure. (A) Supersac intake averaged across drinking test days. Supersac intake in male (B) and female (C) adult rats during each drinking session. A significant difference between males and females in supersac intake averaged across drinking test days within the same adolescent exposure condition is depicted with $, whereas asterisks indicate significant differences in supersac intake between water- and AIE-exposed animals evident at a certain test day (p < 0.05).

**Figure 4.**
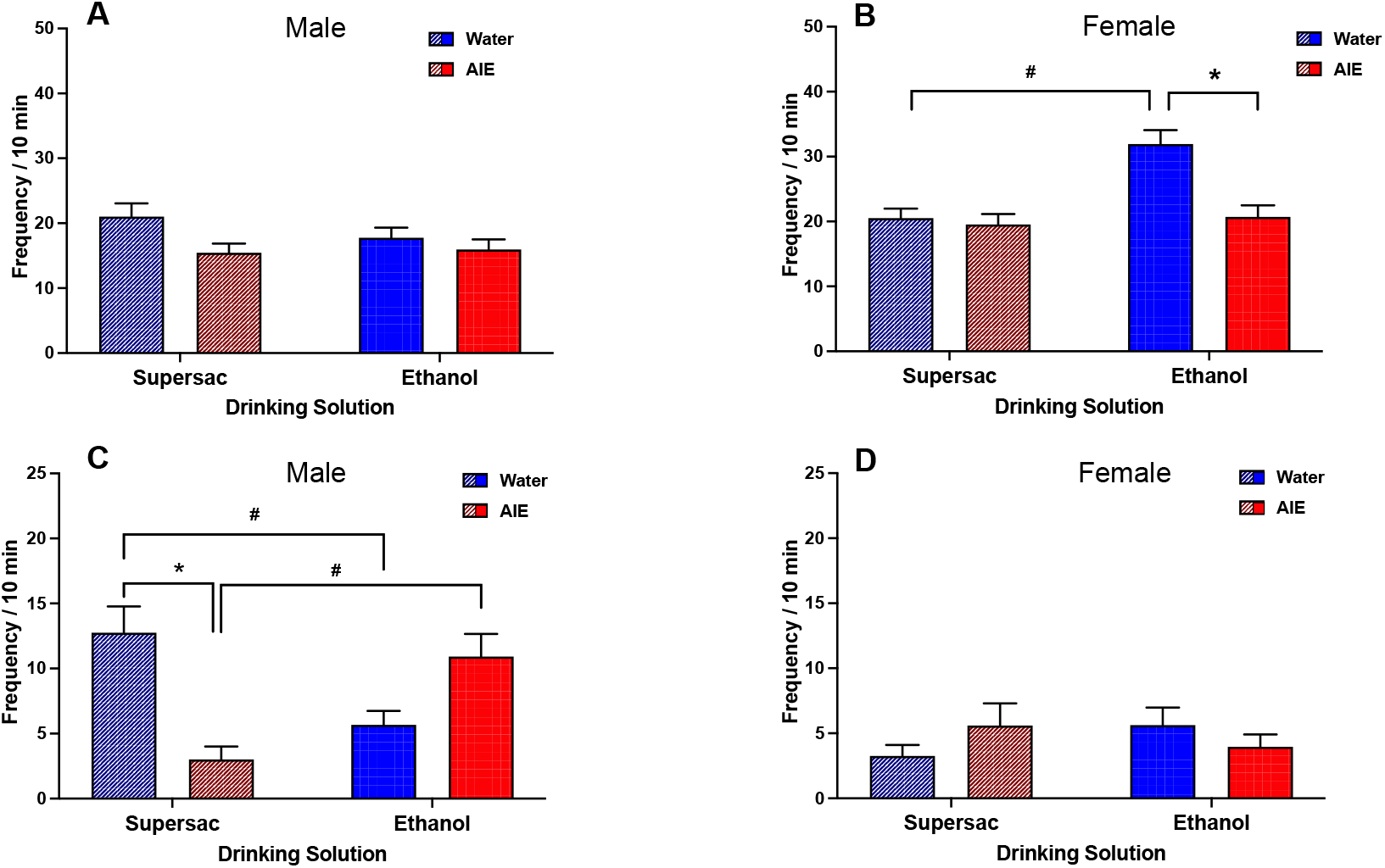
Social investigation (A, B) and play flighting (C, D) demonstrated by male and female rats with the history of water or AIE exposure during social drinking of either supersac only or 10% ethanol solution in supersac. (*) indicates a significant difference between water- and AIE-exposed animals drinking the same solution, whereas (#) denotes differences between animals within the same adolescent exposure condition (water or AIE) drinking different solutions under social circumstances (p <0.05).

### Corticosterone

The ANOVA of corticosterone levels assessed on the last drinking day revealed a main effect of sex, F (1, 165) = 166.7, p < 0.0001, with females demonstrating substantially higher corticosterone levels (overall mean = 62.23 ± 26.67 μg/dL) than their male counterparts (overall mean = 22.32 ± 10.08 μg/dL). However, corticosterone levels were not affected by either adolescent exposure or drinking solution.

### Social Interaction

Social investigation differed as a function of sex, adolescent exposure, and drinking solution, as evidenced by a significant three-way interaction, F (1, 184) = 8.249, p < 0.01. Therefore, separate for each sex 2-way ANOVAs were performed. In males, social investigation during drinking session was not affected by adolescent exposure or drinking solution (see Figure 4A). In females, the 2-way ANOVA of social investigation showed main effects of adolescent exposure, F (1, 92) = 12.6, p < 0.001, and drinking solution, F (1, 92) = 11.78, p < 0.001, as well as adolescent exposure by drinking solution interaction, F (1,92) = 8.363, p < 0.01. Water-exposed females in the ethanol drinking condition demonstrated significantly higher frequency of social investigation than their counterparts ingesting supersac and AIE females drinking ethanol (Figure 4B). A three-way ANOVA of play fighting also revealed a significant interaction of sex, adolescent exposure, and drinking solution, F (1, 184) = 22.96, p < 0.0001. In males, a two-way ANOVA showed a significant interaction of adolescent exposure and adult drinking solution, F (1, 92) = 24.19, p < 0.0001. Post hoc comparisons revealed that AIE males drinking supersac demonstrated significantly less play fighting than their water-exposed counterparts as well as AIE males drinking ethanol, whereas water-exposed males showed less play fighting while drinking ethanol than their counterparts in the supersac drinking condition (Figure 4C). In females, play fighting did not differ as a function of either adolescent exposure or drinking solution (Figure 4D).

### Gene Expression

In males, hypothalamic gene expression was not affected by either adolescent exposure or drinking solution for OXT, OXTR, AVP, and AVPR1a (all ps > 0.05, Figure 5A-D). AVPR1b gene expression differed as a function of drinking solution, F (1, 24) = 14.33, p <0.001 (see Figure 5E), with significantly higher AVPR1b expression evident in males drinking ethanol than those consuming supersac.

**Figure 5.**
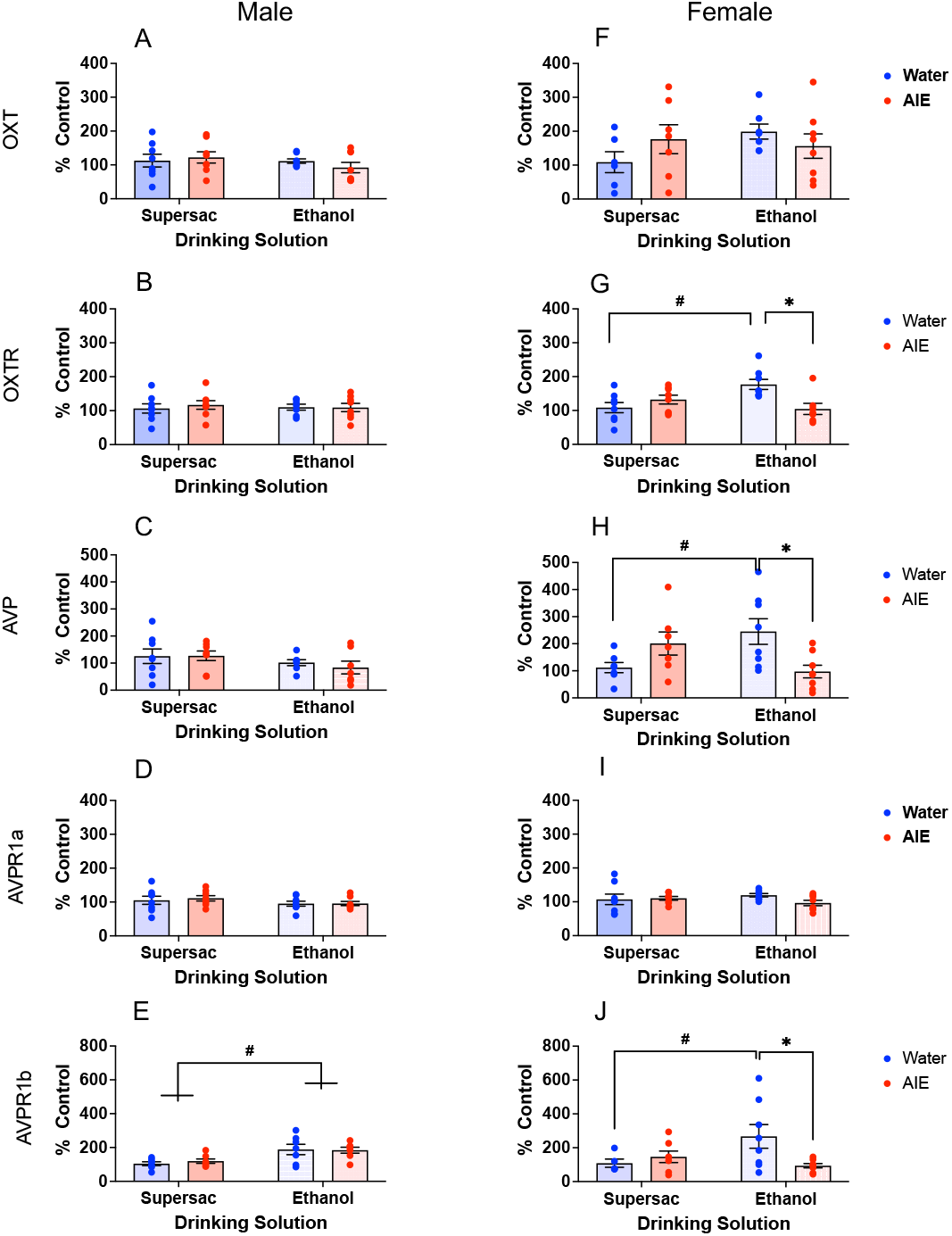
Oxytocin (OXT), oxytocin receptor (OXTR), vasopressin (AVP), vasopressin receptors 1a (AVPR1a) and 1b (AVPR1b) gene expression in the hypothalamus of male (A, B, C, D, E) and female (F, G, H, I, J) rats with the history of water or AIE exposure following the last social drinking test with either supersac only or 10% ethanol in supersac. Data were calculated as a relative change in gene expression using the 2−ΔΔC(t) method, with Cyclophilin A used as a reference gene and the water-exposed animals drinking supersac serving as the ultimate control group within each sex. (*) indicates a significant difference between water and AIE exposure conditions, whereas (#) denotes a significant difference (p < 0.05) in gene expression associated with ethanol versus supersac drinking either within the same adolescent exposure condition (G, H, J) or collapsed across adolescent exposure (E).

In females, neither adolescent exposure, nor drinking solution affected hypothalamic OXT and AVPR1a gene expression (ps >0.05, Figure 5F, I). However, significant adolescent exposure by drinking solution interactions were evident for OXTR, F (1, 27) = 10.41, p <0.01, AVP (1, 26) =11.02, p <0.01, and AVPR1b, F (1, 24) = 5.28, p < 0.05, gene expression (see Figure 5G, H, J). For these genes of interest, water-exposed females drinking ethanol demonstrated significantly higher gene expression than their counterparts drinking supersac as well as AIE females in the ethanol drinking condition.

In general, the genes of interest in the LS were not affected by adolescent exposure and drinking solution (*data not shown*). The only exception was AVPR1a expression that was significantly higher in females drinking ethanol solution (148.5%) relative to those drinking supersac (116%), as evidenced by a main effect of drinking solution, F (1, 28) = 5.18, p <0.05.

## Discussion

This study was designed to test whether AIE-associated alterations of social drinking were sex-specific, with AIE exposure enhancing drinking in males but not females. In general, sex differences in ethanol intake and BECs were evident in water-exposed controls, with females drinking more and achieving higher BECs than males, with these sex differences being eliminated by AIE. During the first two drinking sessions, a substantial reduction in ethanol intake was observed in both sexes exposed to AIE, with AIE-exposed males and females consuming substantially less ethanol than controls. One possible explanation of this drastic suppression of ethanol intake during initial drinking sessions is that novelty of the test situation, being more anxiety-provoking for AIE animals than for their water-exposed counterparts, suppressed ethanol intake in these subjects. We have shown that AIE enhances non-social anxiety-like responding regardless of sex in rats (Varlinskaya et al., 2020), whereas anxiogenic effects of AIE are evident in the novelty-induced hypophagia test of anxiety in male mice under basal conditions (Jury, DiBerto, Kash, & Holmes, 2017) and in AIE-exposed female mice following acute restraint stress (Kasten, Holmgren, Lerner, & Wills, 2021; Kasten et al., 2020). However, it is unlikely that suppression of ethanol intake during initial drinking sessions in AIE males and females was driven by novelty, since no differences in supersac (a palatable sweet solution used as a vehicle for ethanol) intake were evident between AIE- and water-exposed animals at initial testing. An alternative explanation is that during AIE exposure, experimental subjects associated ethanol odor with adverse consequences of either gavage procedure or ethanol intoxication, and ethanol odor became aversive to these animals.

Following the first two drinking sessions, males exposed to AIE and their water-exposed counterparts consumed equivalent amounts of ethanol. During the second 4-day drinking cycle, AIE-exposed males consistently showed greater ethanol intake than water-exposed counterparts on a day to day basis, suggesting that experience with ethanol is necessary for AIE-associated enhancement of ethanol intake to become evident. The observed increase in ethanol intake following AIE in males is in agreement with previous studies that have shown AIE-associated enhancement of ethanol intake in male rats (reviewed in Towner & Varlinskaya, 2020). It is important to note that AIE did not lead to alterations in the intake of supersac, suggesting the elevated intake observed was likely more attributable to ethanol rather than the palatable vehicle. Similar to our findings, previous studies have found no difference in the motivation for acquiring and consuming sweet solutions/palatable food following AIE exposure of male rats (Risher et al., 2013; Towner & Spear, 2021; Varlinskaya et al., 2020; Wukitsch, Moser, Brase, Kiefer, & Cain, 2020).

In addition, we found water-exposed males to display greater fluctuations in intake across individual test days, a pattern potentially representing binge-like drinking and similar to our previous findings with voluntary ethanol consumption (Hosová & Spear, 2017). This pattern of drinking may have been a result of negative consequences associated with the high intake that commonly curb future consumption. In contrast to water exposure, AIE-exposed males showed a more stable drinking over time. The lack of substantial changes in intake over days may be associated with drinking of a certain amount in order to achieve certain BEC levels at which AIE males could experience some desired effects of ethanol without experiencing negative consequences. Furthermore, previous studies have found that AIE reduces the sensitivity to the aversive properties of ethanol (Diaz-Granados & Graham, 2007; Graham & Diaz-Granados, 2006; Saalfield & Spear, 2015; Williams, Nickel, & Bielak, 2018), potentially allowing AIE males to sustain consistent levels of ethanol intake.

In contrast, AIE-exposed females ingested less ethanol than water-exposed controls. The results of previous studies that assessed AIE effects on ethanol intake in females are mixed (for references see Towner & Varlinskaya, 2020), with some reporting AIE-associated decreases in ethanol consumption (Siciliano & Smith, 2001), no effects of AIE (Jury et al., 2017; Rodd-Henrick et al., 2002; Varlinskaya et al., 2017), or increases in ethanol intake (Amodeo et al., 2018; Jacobsen, Buisman-Pijlman, Mustafa, Rice, & Hutchinson, 2018; Maldonado-Devincci, Alipour, Michael, & Kirstein, 2010; Maldonado-Devincci & Kirstein, 2020; Strong et al., 2010). Given that males are predominantly used in ethanol research, more work is needed to more thoroughly characterize the consequences of AIE among females on ethanol consumption in adulthood. It seems unlikely that decreased ethanol intake in females is related to AIE-induced anhedonia, since, intake of supersac was elevated in AIE-exposed females, suggesting that the lack of preference for the sweet taste of ethanol solution did not contribute to the lower ethanol intake. Alternatively, it is possible that the social drinking paradigm was more anxiety-provoking for AIE females, however no differences in the stress hormone CORT were noted between the two adolescent exposure conditions. It is important to consider habituation of the CORT response, since it was measured after the final drinking session thus only providing limited information.

In our previous study, we found no differences in social drinking between exposure conditions within each sex (Varlinskaya et al., 2017). The discrepancy between the current findings and our previous work (Varlinskaya et al., 2017) may be associated with procedural differences between the two studies. Varlinskaya et al. (2017) used a lower dose of ethanol (3.5 g/kg) than the dose used in the current study (4 g/kg) for the AIE procedure. It is likely that the dose of 4 g/kg resulted in higher sustained BECs, thus potentially producing greater alterations than the 3.5 g/kg ethanol dose. Indeed, previous findings suggest more substantial behavioral alterations following higher doses of ethanol during adolescent ethanol exposure (Broadwater, Varlinskaya, & Spear, 2011; Maldonado-Devincci et al., 2010; Maldonado-Devincci & Kirstein, 2020; Matthews, Tinsley, Diaz-Granados, Tokunaga, & Silvers, 2008). Additionally, in Varlinskaya et al. (2017) study, experimental subjects were housed with non-littermates from weaning, through AIE and the drinking procedure, whereas animals were housed with littermates in the present study. It is possible that the stress of re-housing in early life mitigated some of the differences between AIE- and water-exposed animals that were evident in the current study.

AIE has been shown to elicit social alterations in males, but not females (Dannenhoffer et al., 2018; Varlinskaya et al., 2014; Varlinskaya et al., 2017; Varlinskaya et al., 2020), with some of these alterations reversed by acute ethanol in a low dose range (Varlinskaya et al., 2014). In addition, social drinking has been shown to increase social play behavior in AIE males, but not water-exposed controls (Varlinskaya et al., 2017). As in this previous study, ethanol consumption increased play fighting only in AIE males. In contrast, water-exposed males exhibited a reduction in social play relative to their counterparts consuming supersac, a behavioral response representing ethanol-induced social inhibition. Social investigation was not affected by either adolescent exposure or drinking solution in males. Taken together with ethanol intake data, this finding suggests that AIE males may moderate their ethanol intake to sustain BECs at a certain level that facilitates play fighting.

Water-exposed females demonstrated greater social investigation while drinking ethanol relative to their supersac drinking counterparts and AIE-exposed ethanol drinking females. This ethanol-induced facilitation of social investigation in control females was rather surprising, especially given that an acute challenge with 0.5 g/kg ethanol increased social investigation in females regardless of adolescent exposure condition (Varlinskaya et al.,2014). Interestingly, BECs achieved by water-exposed females were around 40 - 60 mg/dl, whereas in the Varlinskaya et al. (2014) study, water- and AIE-exposed females showed ethanol-induced facilitation of social investigation following systemic ethanol administration at substantially lower BECs (around 20.0 mg/dl). At BECs higher than 40 mg/dl, water- and AIE-exposed females showed no changes in social investigation following acute ethanol challenge (Varlinskaya et al., 2014). Together, these results suggest changes in sensitivity to socially facilitating effects of ethanol in water-but not AIE-exposed females. These changes in sensitivity may be associated with repeated exposure to ethanol during social drinking sessions, with water-exposed females demonstrating chronic tolerance to the socially facilitating effects of ethanol and showing this facilitation at substantially higher BECs.

Following the final drinking session, we assessed changes in OXT and AVP neuropeptide systems gene expression. These sexually dimorphic brain systems (Dumais & Veenema, 2016) are important modulators of various social behaviors (Caldwell, 2017; Harper et al., 2019; Johnson & Young, 2017; Veneema & Neumann, 2008; Zoicas et al., 2014). In addition, recent evidence suggests that these systems are affected by ethanol (Allen et al., 2016; Dannenhoffer et al., 2018; Peters et al., 2017; Przybycien-Szymanska et al., 2010; Rivier & Lee, 1996; Silva et al., 2002; Silva et al., 2004) and might contribute to voluntary ethanol consumption (Edwards et al., 2012; MacFadyen et al., 2016; Zhou et al., 2017). In males, adolescent exposure did not affect gene expression. The only change evident was an increase in hypothalamic AVPR1b gene expression in males drinking ethanol relative to those drinking supersac. In contrast, water-exposed females drinking ethanol had higher AVP and AVPR1b gene expression than their counterparts drinking supersac, However, the only group that did not show changes in AVP and AVPR1b mRNA levels while drinking ethanol were AIE-exposed females, suggesting that adolescent ethanol exposure made these females insensitive to the effects of ingested ethanol on the AVP/AVPR1b system. It seems that the observed changes in the hypothalamic AVP system (i.e., increases in AVP and/or AVPR1b gene expression) were associated with acute effects of ingested ethanol rather than consequences of multiple drinking episodes, since repeated exposure to ethanol has been shown to suppress AVP gene expression in the hypothalamus of adult rodents (Ishizawa, Dave, Liu, Tabakoff, & Hoffman, 1990; Madeira et al., 1997; Sanna et al., 1993), More work is needed to determine relationships between the AVP/AVPR1b system, social drinking, sex, and prior experience with ethanol.

We also found increases in hypothalamic OXT and OXTR gene expression in water-exposed females drinking ethanol. To our best knowledge, a relationship between OXT gene expression and ethanol intake in females has not been investigated, although acute OXT has been shown to reduce ethanol intake in both sexes (Caruso, Robins, Fulenwider, & Ryabinin, 2021; King & Becker, 2019; Peters et al., 2017). Furthermore, evidence suggests that OXTR knockout leads to increases in ethanol consumption (Rodriguez, Smith, & Caldwell, 2020), suggesting that the observed activation of the OXT system was a compensatory mechanism of curbing ethanol intake that was relatively high in water-exposed females. Interestingly, the role of OXTR in the modulation of ethanol intake may be sex-specific, given that OXTR knockout increased intake in females but had no effect in males (Rodriguez et al., 2020). There was no ethanol-induced changes in the OXT system gene expression in AIE-exposed females, which may be related to relatively low ethanol intake in these animals. An alternative explanation is that the activation of the OXT in water-exposed females drinking ethanol was associated with social behavior. Indeed, water-exposed females were the only group socially facilitated by ethanol, as indexed by increases in social investigation during the final social drinking episode. Future work should assess whether drinking alone can elicit similar increases in OXT/OXTR gene expression in adult females.

Importantly, the observed changes in the OXT/AVP systems were evident in water-exposed females which displayed the highest level of ethanol intake and was associated with enhanced social investigation. It is possible that under normal circumstances, activation of these neuropeptide systems in the HYPO is an important contributor to ethanol-induced changes in social behavior in females, but not males. It is likely that this neuropeptide modulation of the facilitating effects of ethanol on social investigation is altered by exposure of adolescent females to ethanol. However, it should be clarified whether the observed changes in gene expression are specific to a social drinking paradigm or if similar alterations can be seen after an acute ethanol challenge.

Overall, the current study found that adolescent ethanol exposure produced sex-specific alterations in social drinking, with AIE moderately increasing ethanol consumption in males and decreasing it in females. AIE-exposed males demonstrated increases in play fighting and water-exposed males showed decreases in social investigation during the final drinking session. In contrast to their male counterparts, water-exposed females demonstrated increases in social investigation during the final drinking session. These behavioral responses to ingested ethanol were accompanied by alterations in OXTR, AVP, and AVPR1b gene expression in the hypothalamus, most predominantly among water-exposed females. Together, these results provide novel information regarding sex-dependent alterations associated with AIE and suggest sex-specific effects of ingested ethanol on social investigation and the hypothalamic OXT/AVP system. More work is needed for better understanding the contribution of these two neuropeptide systems in the sex-specific drinking and social behavior patterns observed.

## Acknowledgements

Supported by NIH grants U01 AA019972, T32 AA025606 and F31 AA029300. Any opinions, findings, and conclusions or recommendations expressed in this material are those of the author(s) and do not necessarily reflect the views of the above stated funding agencies. The authors have no conflicts of interest to declare.

